# Influence of color and brightness on ontogenetic shelter preference by prawns (*Macrobrachium rosenbergii*)

**DOI:** 10.1101/2020.11.05.368035

**Authors:** Felipe P. da Costa, Maria F. Arruda, Karina Ribeiro, Daniel M. A. Pessoa

**Author notes:** Corresponding author: Caixa Postal 1511, 59078-970 Natal, Brazil.

## Abstract

The giant river prawn (*Macrobrachium rosenbergii*), native to rivers and river mouths of different Asian countries, is a heavily widespread species that has been introduced around the world due to its great commercial importance. These prawns are farmed under many different conditions that might translate to a great range of light environments, which impact their behavior and productivity. Here, as a contribution for prawns’ welfare and economical productivity, we present the first study employing both visual modeling and behavioral data to evaluate ontogenetic changes on color preference of juveniles and adults of *M. rosenbergii*. For this purpose, we offered ten shelters of different colors to juveniles and adults and registered their preference. Our results showed that the criterion for shelter preference changed with ontogeny, since juveniles chose shelters based on chromaticity (preference for blue), while adults based their decisions on brightness (preference for dark gray). This preference of adults for dark colors is probably associated with a light avoidance behavior. We recommend providing blue shelters for juveniles and dark shelters for adults.

## 1. Introduction

The giant river prawn *Macrobrachium rosenbergii* (De Man, 1879) is native to a region that encompasses Malaysia, East India, West Indonesia, Gulf of Bengal, and Gulf of Thailand (Holthuis & Ng, 2010). In their natural environment, the larvae inhabit estuarine environments (Sandifer & Smith, 1985), are planktonic (New, 2002) and go through 11 larval stages (from Zoea I to Zoea XI) (Sandifer & Smith, 1985). After a larval life of 20 to 50 days, the larvae undergo metamorphosis into post-larvae (Sandifer & Smith, 1985), which are benthic and begin migration to freshwater environments, where they remain until adulthood (New, 2002) and mate (Sandifer & Smith, 1985). Then, the ovigerous female migrates to estuarine environments, where the larvae hatch from the eggs, restarting the cycle (Sandifer & Smith, 1985).

*M. rosenbergii* is a species of great commercial importance (Engle, Quagrainie, & Dey, 2016; Zeng, Cheng, Lucas, & Southgate, 2012) that, in 2018 alone, accounted for the production of 234,400 tons of food worldwide (FAO, 2020). These prawns are farmed under various conditions (Coyle, Alston, & Sampaio, 2010; Daniels, Cavalli, & Smullen, 2010; Valenti, Daniels, New, & Correia, 2010; Valenti, New, Salin, & Ye, 2010), that might translate to a great range of light environments, which have the potential to impact prawns’ behavior and productivity. In fact, one study has already shown that food color affects prawns’ larvae feeding behavior (Yong, Kawamura, Lim, & Gwee, 2018). In different species, color preference has also been related to the selection of appropriate habitats (Gu et al., 2017; Havel & Fuiman, 2017; Strader, Davies, & Matz, 2015). For instance, changes in color preference throughout the ontogenetic development of some caridean shrimps (Lysmatidae) have been related to the physical properties of the environments occupied by them at different stages of development (Johnson & Rhyne, 2015). Therefore, the study of color preference by species of economic interest might exert an important role in animal welfare and food productivity.

Morphologically, the eyes of *M. rosenbergii* change throughout ontogenetic development, since larvae have functionally apposition eyes, whereas adults have functionally reflecting superposition eyes (Nilsson, 1983). The superposition eyes are more efficient for gathering light and can be advantageous in low light conditions, but the transformations necessary for the appearance of a superposition eye may not be complete in post-larvae, as seen in a caridean shrimp (Douglass & Forward, 1989). Regarding the visual sensitivity of the species, it was found that dark-adapted individuals exhibit a light absorption peak at 563 nm, which corresponds to the yellow-red region of the spectrum (Matsuda & Wilder, 2014), although the authors don’t specify how many photoreceptor types would be accounting for the sensitivity curve. Still, in spite of the great deal of experiments that have already been conducted with giant prawns (Chong-Carrillo et al., 2016), only recently their visual system begun to be studied through behavioral experiments (Kawamura, Bagarinao, Yong, Faisal, & Lim, 2018; Kawamura, Bagarinao, Yong, Fen, & Lim, 2017; Kawamura, Bagarinao, Yong, Jeganathan, & Lim, 2016; Kawamura, Yong, Wong, Tuzan, & Lim, 2020).

Through visual modeling studies, a strategy that has currently gained popularity, it is possible to assess which spectral information available in the environment could be exerting an adaptive function. During visual modeling, we infer how a given animal’s visual system is stimulated by observing a particular object under a specific illuminant (Olsson, Lind, & Kelber, 2018). In other words, just by knowing how many kinds of photoreceptors (and their peak sensitivities) there are in an animal’s eye, the spectrum of ambient light, and the color of an object of interest, we can suppose how the object should be seen by that observer. Yet, although variations in the type and number of photoreceptors are usually related to the dimensionality of color vision, only behavioral tests can verify an animals’ color perception (Jacobs, 1996). So, it is important to couple visual modeling data with behavioral experiments that can validate them (Lind & Kelber, 2009).

To our knowledge, availability of chromatic and achromatic cues to *M. rosenbergii* have never been properly analyzed by means of visual modeling. The experiments already carried out either did not try to control the brightness of the stimuli or tried to do so without correctly taking into account the prawns’ visual system. In color preference experiments, the choice of stimuli colors should consider the eye of the beholder (Hill, 2002), not human vision. By using visual modeling, we can control the colors of the stimuli to be presented to prawns in a rigorous manner, as in studies with other animal species (Detto, 2007; Escobar-Camacho et al., 2019; Olsson, Lind, & Kelber, 2015; Siebeck, Wallis, Litherland, Ganeshina, & Vorobyev, 2014).

Therefore, here we developed the first color preference study on *M. rosenbergii* employing visual modeling, as a more rigorous control for stimuli brightness and chromaticity, according to the visual system of prawns. Our aim is to investigate whether their preference for different colors and brightness changes over their development. Since *M. rosenbergii* spontaneously occupies experimental shelters (Santos, Pontes, Campos, & Arruda, 2015), we analyzed the preference that two benthic developmental stages (i.e. juveniles and adults) show for shelters of different chromaticities and brightnesses. Our hypothesis is that *M. rosenbergii* changes their color preference gradually throughout ontogeny, regardless of the farming conditions. This ontogenetic shift would be adaptative in natural populations because the spectral composition (i.e. color) of the surrounding natural photic environment changes as they start migrating from estuaries to freshwater ecosystems. Since this migration only begins when *M. rosenbergii* reaches its post-larvae developmental stage, we predict that adults will be more prone than juveniles to exhibit strong preferences for longer wavelengths, such as yellow and orange, which are colors associated to freshwater bodies (Stoffels, 2013).

## 2. Material and methods

### 2.1 Ethical statement

Our research was approved by the Ethics Committee on The Use of Animals of our institution (protocol 042/2018) and is in accordance with Brazilian law. It complies with ARRIVE guidelines and was carried out in accordance with the U.K. Animals (Scientific Procedures) Act, 1986, and associated guidelines, EU Directive 2010/63/EU for animal experiments.

### 2.2 Animal maintenance

At the Laboratory of Sensory Ecology, of the Federal University of Rio Grande do Norte, we kept 39 juveniles of *Macrobrachium rosenbergii*. Animals were housed in a collective aquarium (100 x 50 cm, 40 cm water column), with transparent water, sandy substrate and aeration. They were subjected to a 12-hour light/12-hour dark light cycle (light from 6:00 a.m. to 6:00 p.m.). Two fluorescent lamps provided a light intensity of approximately 320 lux (measured with an Extech Instruments HD 400 Light Meter), which should be sufficient for color discrimination (Kawamura et al., 2018).

We kept the physical-chemical parameters of the water at optimum levels for the prawns (pH: 7.0-7.5; ammonia: 0 ppt; temperature: 26-28 °C), and fed the animals twice a day (morning and afternoon) with commercial food containing 42% crude protein. Twenty percent of the aquarium’s water was changed twice a week.

### 2.3 Experimental apparatus

We also made available transparent water, sandy substrate and aeration in two experimental aquaria, with aeration positioned, approximately, at the center of each aquarium. In each experimental aquarium, we arranged ten shelters, in two rows of five. We build the shelters with plastic coated paper, folded in an appropriate way. Each aquarium had six gray shelters, each one with a different brightness, besides a blue, a green, a yellow and an orange shelter (Figure 1).

**Figure 1.**
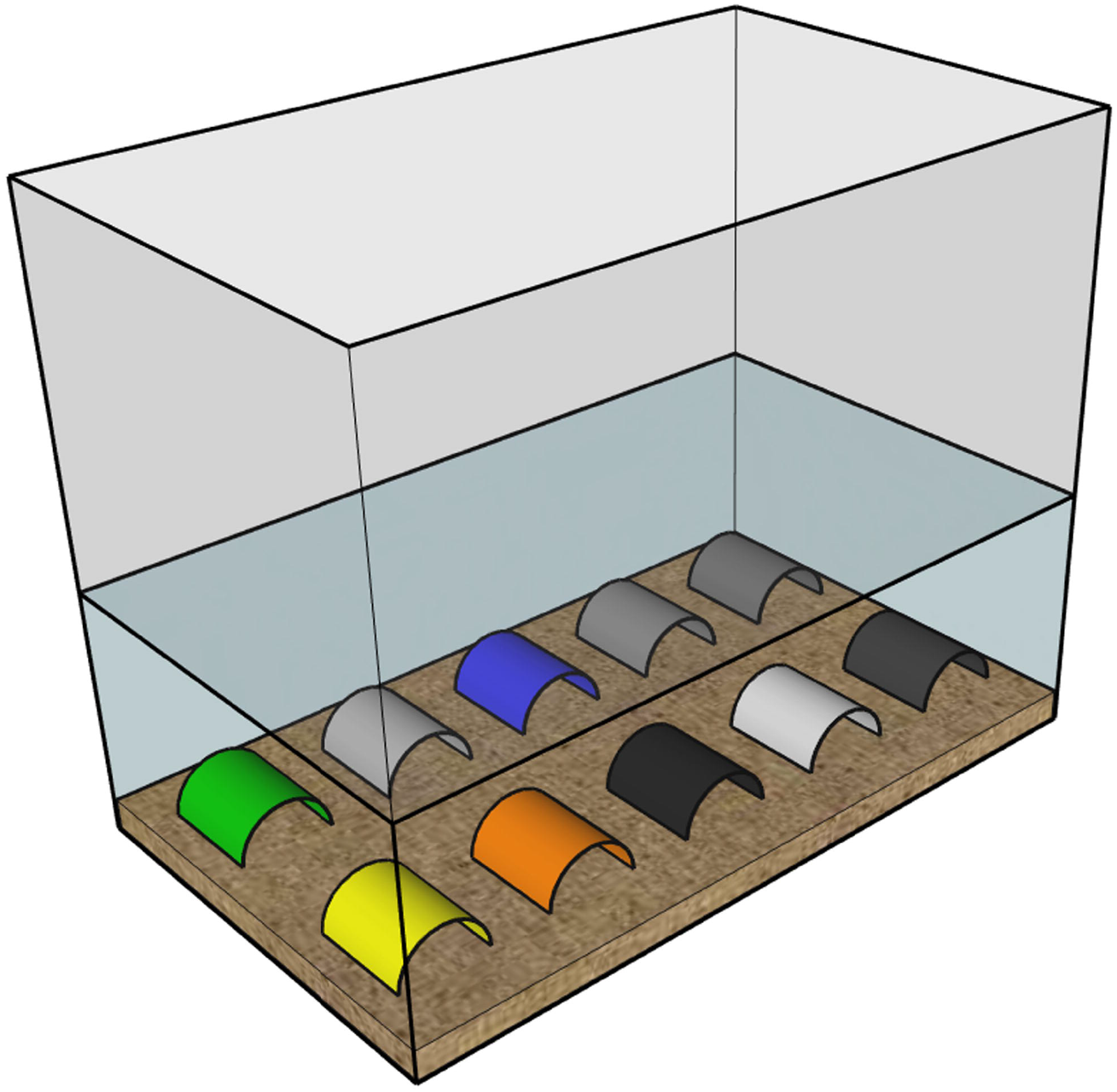
Schematic view of the arrangement of shelters in an experimental aquarium. The prawns were individually tested in aquaria with 10 colored shelters arranged in two rows of five.

### 2.4 Shelters’ colors and brightness

For choosing our stimuli colors, we printed 101 different color patches that were coated with plastic and had their reflectance spectra measured by us with a spectrophotometer (USB4000 UV-VIS Fiber Optic Spectrometer, Ocean Optics, Inc.). We coupled the spectrometer to a bifurcated optical fiber (QR450-7-XSR, Ocean Optics, Inc.), also attached to a light source (DH-2000-BAL, Ocean Optics, Inc.). A white standard surface (WS-1-SL, Ocean Optics, Inc.), and the obstruction of the light source and optical fiber, were used as, respectively, the white and the black standards, for system calibration. We also measured the illuminant of the experimental room with the spectrometer coupled to an optical fiber (QP450-2-XSR, Ocean Optics, Inc.), attached to a cosine corrector (CC-3-UV-S, Ocean Optics, Inc.). We calibrated this spectrometric system with a calibration light source (LS-1-CAL, Ocean Optics, Inc.).

We run visual models, through pavo 2 (Maia, Gruson, Endler, & White, 2019), a package for R 3.4.1 (R Development Core Team, 2020), to infer how each color patch would be seen according to the prawn’s visual system. Our model computed the quantum catches absorbed by each type of photoreceptor described for prawns, when the animals visualized each shelter (reflectance spectra in Figure 2), that was illuminated by the fluorescent lamps of our experimental room (Figure 3). The absorption peaks of the photoreceptors of another prawn species, *Palaemonetes poludosus* (380 nm and 555 nm) (Goldsmith & Fernandez, 1968), were adopted in our model, since data were not available for *M. rosenbergii. P. poludosus* is the taxon most closely related to *M. rosenbergii* for which photoreceptors’ absorption peaks have been established. Using photoreceptors’ absorption peaks of related species is a viable alternative, since small variations in estimated peaks do not carry a great influence in visual modeling results (Lind & Kelber, 2009; Olsson et al., 2018).

**Figure 2.**
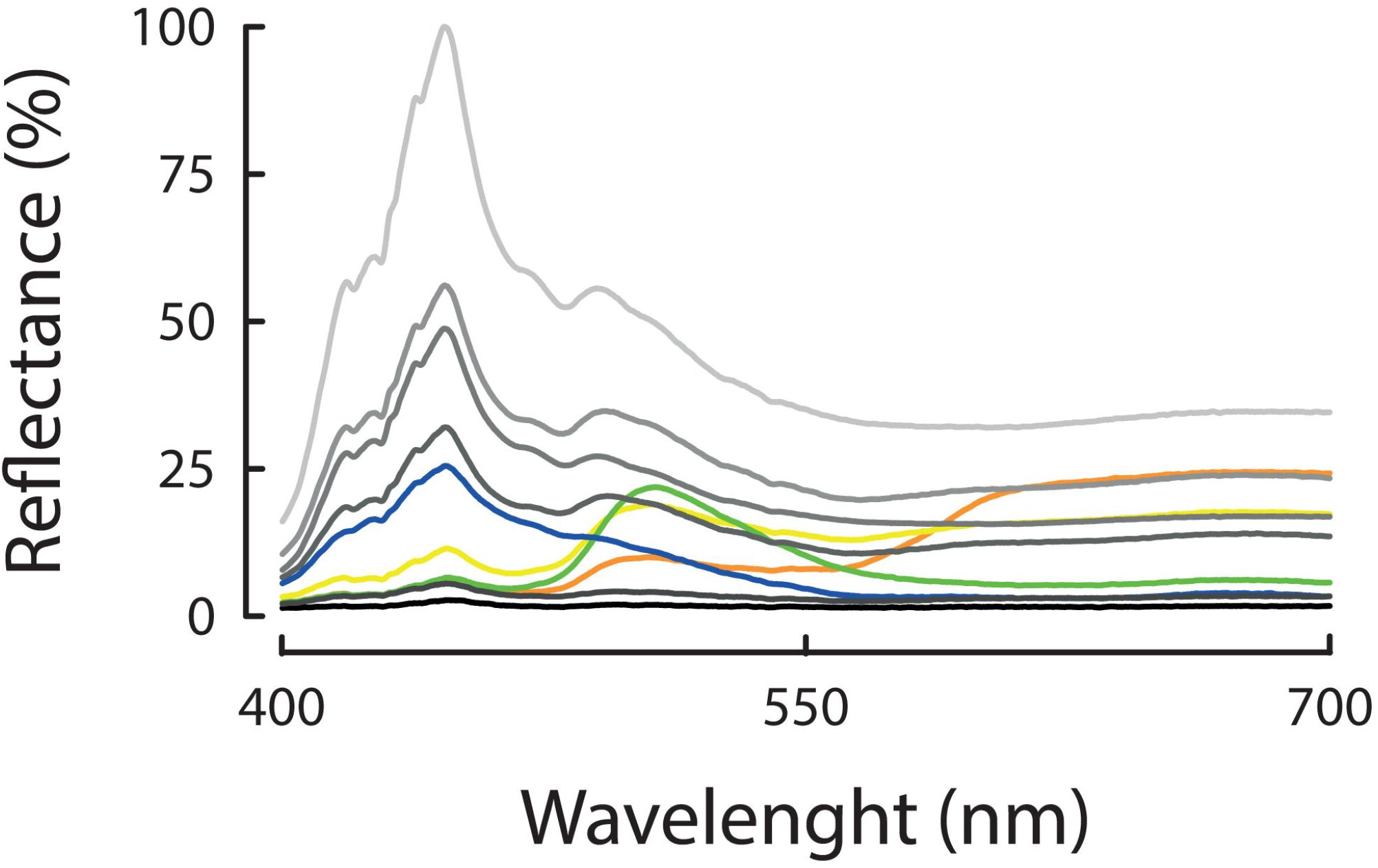
Reflectance spectra of experimental shelters. The reflectance of each curve is represented as a percentage, compared to the maximum reflectance of the most reflective curve. Each curve is represented approximately the same color as the respective shelter.

**Figure 3.**
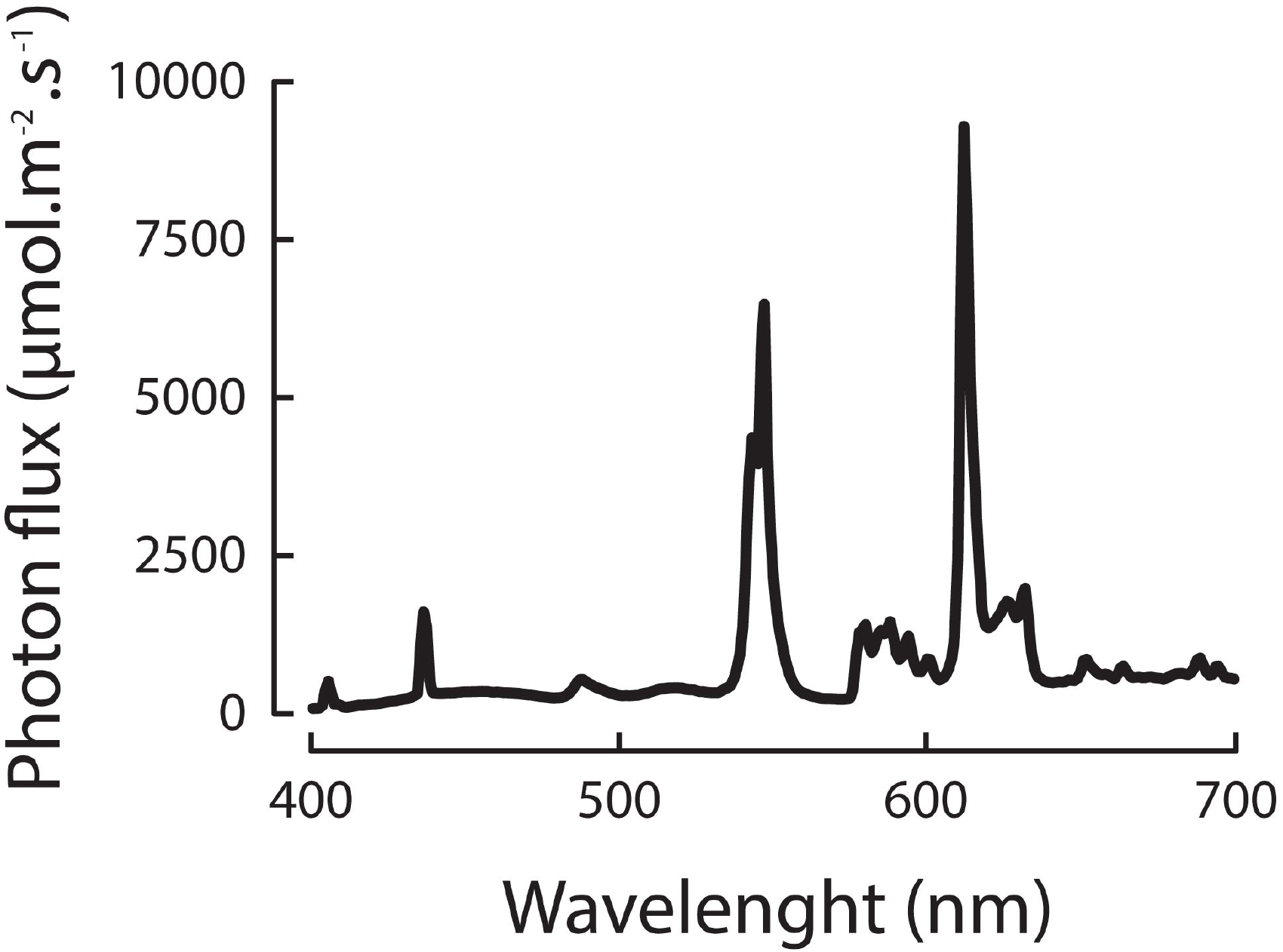
Illuminant spectrum of the experimental room. Illumination was provided by two fluorescent lamps.

In order to calculate the signal of each photoreceptor, we divided the amount of light reflected by a stimulus (shelter) and captured by a specific photoreceptor class (stimulus quantum catch), by the amount of light reflected by a perfect white surface (white standard) and captured by the same photoreceptor class (maximum quantum catch). Following Detto (2007), we computed achromatic signals for each stimulus, as the sum of signals of short wavelength (S signals) and long wavelength (L signals) photoreceptors. For chromatic signals, we divided S signals by L signals (Siebeck et al., 2014). Chromatic and achromatic signals were plotted in a chromaticity-luminance diagram (Figure 4), that show how our stimuli varied with respect to color and brightness. We chose not to use the RNL model (Vorobyev & Osorio, 1998) because it demands visual parameters (Olsson et al., 2018) that are still unknown for prawns.

**Figure 4.**
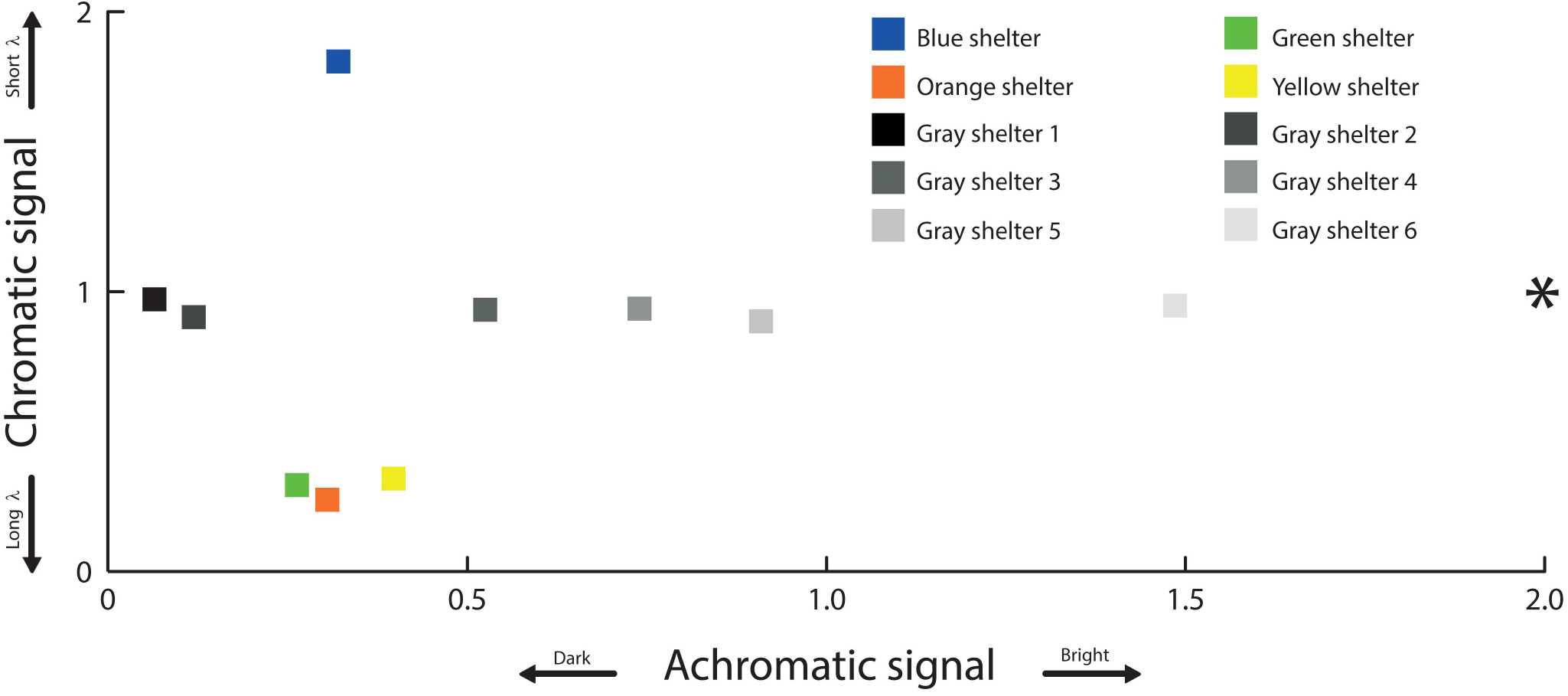
Chromaticity-luminance diagram for the visual system of prawns. Inferred chromatic and achromatic signals, determined for a perfect white surface (asterisk) and for shelters employed in our experiments (squares), are represented. Distance between squares indicate color and brightness difference in chromatic and achromatic signal axes, respectively. Shorter λ = shorter wavelengths (e.g. blues); Longer λ = longer wavelengths (e.g. yellows).

### 2.5 Experimental design

In experiment 1, we individually tested 39 juveniles about two months after metamorphosis (body weight: 0.46 ± 0.13 g; total length: 3.89 ± 0.36 cm), two juveniles per day, each one in a smaller experimental aquarium (30 x 50 cm). The shelters, in experiment 1, measured 7 cm depth x 6 cm width x 3 cm height. The position of the different shelters for each animal was randomized.

In experiment 2, we individually tested 24 of the 39 previously tested individuals, when they were already adults about seven months after metamorphosis (body weight: 6.4 ± 3.2 g; total length: 8.49 ± 1.84 cm), one adult prawn per day, in a larger experimental aquarium (50 x 100 cm). The shelters, in experiment 2, measured 7 cm depth x 9 cm width x 5 cm height. Otherwise, the two experiments were identical. Differences in sample size between experiments 1 and 2 were due to animal mortality.

In both experiments, we fed all prawns in the collective aquaria at 7:30 a.m., removed each prawn, that was about to be tested, from the collective aquarium and transferred it to the center of an experimental aquarium at 8:00 a.m. From 9:00 a.m. to 5:30 p.m, at every 30 min, we recorded if the animal was inside/on top of a shelter (shelter occupancy), or if it was away from any shelter (no choice), following Kawamura et al. (2017). After the experimental session, we removed the prawn from the experimental aquarium, recorded its weight and total length, and transferred it to another collective aquarium, in which individuals that had been already tested were kept. All animals were fed again at the end of the day.

### 2.6 Statistical analysis

We performed ten chi-square tests, one for each shelter, for unequal expected proportions, to verify if the occupancy of each shelter differed from the expected 10% of all cases. We employed the Bonferroni correction to account for multiple comparisons and set our α to 0.005. All tests were performed in BioEstat 5.3 (Ayres & Ayres Jr, 2007).

## 3. Results

### 3.1 Visual modeling

Regarding our stimuli, the blue shelter generated a chromatic signal that stood out from green, yellow and orange shelters (Figure 4), a strong indication that the blue shelter could be perceived, by the prawns’ visual system, as being of a different color. In contrast, these four colored stimuli (i.e. blue, green, yellow, and orange shelters) only showed discrete variations of achromatic signals, with the yellow shelter being slightly lighter than the others (Figure 4). Moreover, all these colored stimuli, according to the visual system of the prawns, were of intermediate brightness when compared to gray shelters. That is, in the absence of chromatic information, animals should not be able to tell colored and gray shelters apart, and would have a great range of achromatic options, including darker (gray shelters), intermediate (gray and colored shelters) and brighter (gray shelters) stimuli, from which they could choose.

### 3.2 Shelter preference

The records of juveniles and adults occupying the shelters are shown in Figure 5. Juveniles occupied the bluest available shelter (i.e. the one with the highest chromatic signal) significantly more than expected by chance (*p* = 0.0034), while occupied all the other available shelters (i.e. those with intermediate and low chromatic signals) at expectancy levels (*p* ≥ 0.009). Adults, on the other hand, occupied two of the most darker available shelters (i.e. those ones with the lowest achromatic signals), gray 1 (*p* < 0.0001) and gray 2 (*p* = 0.0001), significantly more than expected by chance, while occupied the most brighter available shelter (i.e. the one with the highest achromatic signal), gray 6 (*p* = 0.0010), significantly less than expected. Adults occupied all other shelters (i.e. those with more intermediate achromatic signals) at expectancy levels (*p* ≥ 0.0435).

**Figure 5.**
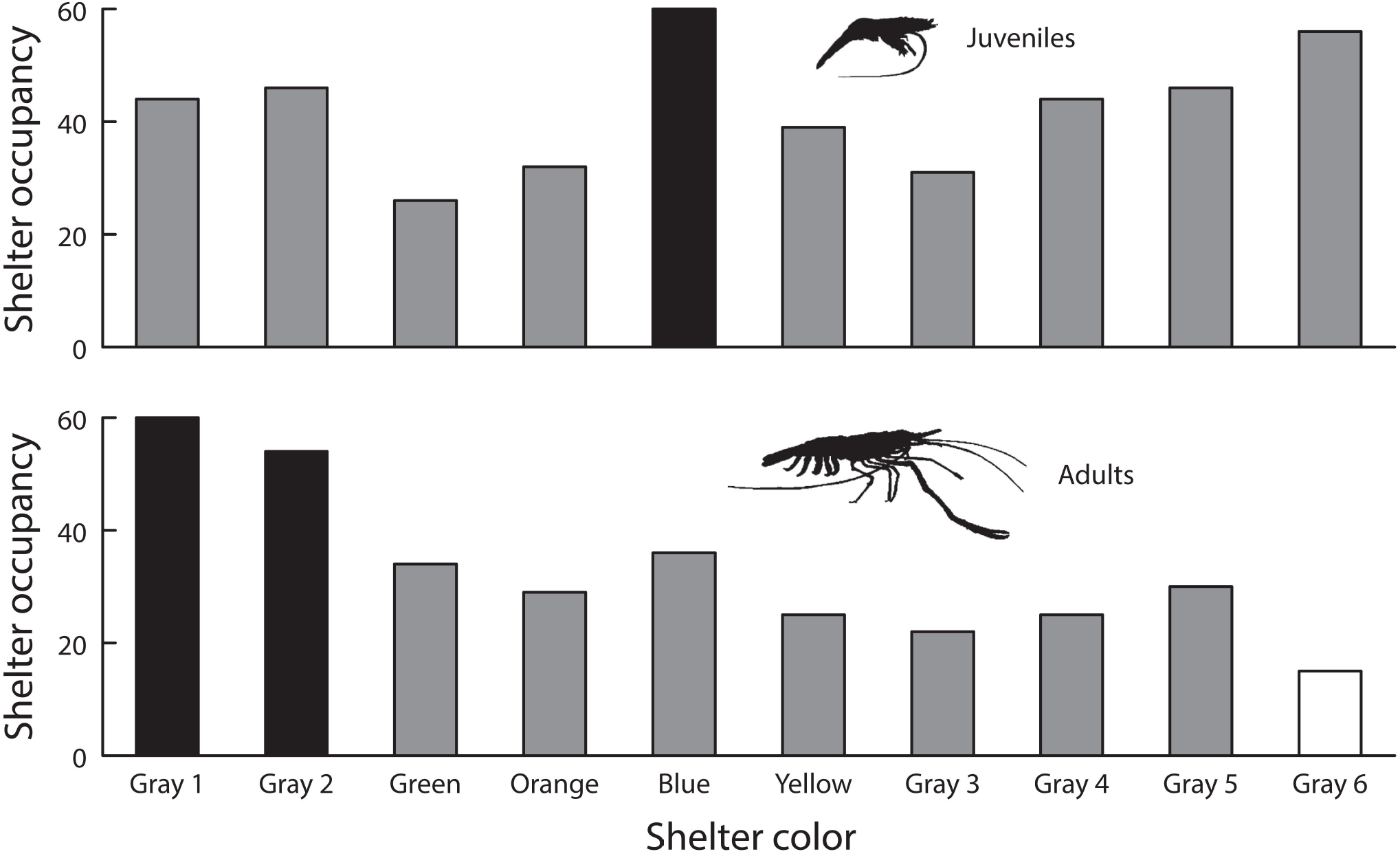
Preference for shelters of different colors by juveniles and adults of giant river prawns (*M. rosenbergii*). The bars represent the number of records of occupancy for each shelter. Black bars represent shelters that were chosen significantly above expectancy level. Gray bars represent shelters chosen at expected frequencies. The white bar represents a shelter chosen significantly below expectancy levels.

## 4. Discussion

Our study is the first to employ visual modeling to study color preference in *Macrobrachium rosenbergii*. The computation of quantum catches, for prawn’s visual system, enabled us to choose blue, green, yellow and orange stimuli, that satisfied two criteria: 1) differed chromatically from the gray shelters, presenting a very high or a very low chromatic signal; and 2) exhibited more intermediate brightness levels, when compared to gray stimuli. Our visual model showed that, for the prawns, the blue shelter presented a chromaticity very different from the chromaticity of the green, the yellow and the orange shelters. On the other hand, the green, the yellow and the orange shelters were very similar in chromaticity for the prawns, probably due to a dichromatic color vision maintained by the interaction of two different types of photoreceptors (Jacobs, 1996). Nevertheless, we must emphasize that our results should be interpreted with caution, since it is not already established if the visual system of *M. rosenbergii* works through chromatic opposition.

Our prediction that adult subjects would prefer longer wavelength colors (i.e. yellow or orange shelters) was not corroborated, since juveniles preferred blue shelters, while adults preferred darker ones. In our study, the stimulus that caught juvenile’s attention (i.e. the blue shelter) was similar in brightness, and chromatically different, from other colored stimuli (i.e. green, yellow, and orange shelters), which were disfavored. This is a strong indication that the blue preference shown by juveniles was based on color alone, not brightness, which is the first indication of color vision in these animals. Adult preference, on the other hand, was probably based on brightness, since different neutral stimuli (gray shelters) were very similar in chromaticity and very different in brightness. Still, our results are not enough to determine whether adult *M. rosenbergii* enjoy color vision, since there is a possibility that they prefer darker shelters despite of a color vision sense. In fact, it has already been reported that adults favor the occupancy of shaded areas of the pond (Karplus & Harpaz, 1990) and that the use of shelters by this species might be linked to a light avoidance behavior (Costa & Arruda, 2016). So, the preference for darker shelters by adults, in our study, may be explained by a tendency to avoid light expressed by the species, which should facilitate their escape from predators (Dabbagh & Kamrani, 2011). This preference for darker shelters has also been observed in other species of the genus *Macrobrachium* (Balasundaram, Jeyachitra, & Balamurugan, 2004; Mariappan & Balasundaram, 2003).

The ontogenetic changes in color preference exhibited by our animals, from blue to dark gray, are consistent with what has been found in less controlled experiments performed by other laboratories (Juarez, Holtschmit, Salmeron, & Smith, 1987; Kawamura et al., 2017; Kawamura et al., 2016; Kawamura et al., 2018; Yong et al., 2018), however, our data demonstrate that these shifts in color preference seem to occur later in life. Regarding the preference of juveniles, our findings are in line with what has been shown for the color preference of the larvae of *M. rosenbergii* in studies employing different colors of beads (Kawamura et al., 2016) and feed (Yong et al., 2018). Similarly, the preference for darker stimuli by our adult subjects is in line with what has been verified in previous studies that tested post-larvae of *M. rosenbergii* (Kawamura et al., 2017), which were shown to prefer black shelters over blue or green ones, and juveniles (Juarez et al., 1987), that preferred black over blue or white substrates. Kawamura et al. (2020), on the other hand, report that the yellow background was the most preferred by larvae and was avoided by post-larvae. The authors also claim that larvae preferred brighter backgrounds, while post-larvae preferred darker ones. However, these authors were unable to satisfactorily consider the animal’s own visual system, since nothing was done to assess the chromaticity of the different stimuli.

The discrepancies found between our results and previous studies might result from the lack of precise control over stimuli and also from methodological differences. For instance, Kawamura et al. (2017), Kawamura et al. (2020) and Juarez et al. (1987) tested *M. rosenbergii* in groups, while we adopted individual testing. We know these animals are aggressive from a young age (Silva & Arruda, 2015), and since Kawamura et al. (2017) reported agonistic interactions among their experimental individuals, we can′t rule out the possibility that their results were biased by hierarchy. For example, in a group context, it is plausible to imagine that a prawn might have avoided occupying a preferred spot that was already occupied by a more dominant individual, forcing it to opt for a suboptimal choice.

Hierarchy can even influence the expression of vision-related genes. Aziz, Rahi, Hurwood and Mather (2018) compared gene expression of the three morphotypes found in populations of *M. rosenbergii* and found higher expression levels of long wavelength opsin genes in the eyestalk and hepatopancreas of small males, which are the most subordinate morphotype. Hence, they suggest that small males could present a better color discrimination, which would help avoiding dominant and subdominant morphotypes, that carry blue and orange claws, respectively. In addition, Santos et al. (2015) noticed a difference in color preference between sex, verifying that females preferred red and orange shelters, while males preferred black ones. Since we did not sex our prawns, and neither did Juarez et al. (1987), Kawamura et al. (2017) or Kawamura et al. (2020), we should not rule out sex bias as a possible explanation for the ontogenetic discrepancies found between studies. Additional behavioral studies should be encouraged, with careful control of stimuli’s chromaticity and brightness, in order to study the development of a preference for chromatic and achromatic cues in males and females of *M. rosenbergii* and to elucidate whether adults have color vision.

In conclusion, our results strongly indicate that *M. rosenbergii* uses chromaticity alone to distinguish between stimuli. Juveniles show a higher preference for blue shelters, which is based on chromaticity, since, according to our visual model, other shelters with comparable brightness were not preferred by the animals in the same way. Later in life, prawns direct their preference to darker shelters, and avoid brighter ones, which seems to be based, mostly, on brightness cues. We suggest that farmers should provide blue shelters for juveniles and dark shelters for adults, to favor shelter occupancy, which can improve productivity and animal welfare.

## Acknowledgments

We would like to thank Felipe N. Castro, for helpful statistical advices; Beatriz A. de Souza, Charles S. Torres, Eider R. de Farias Filho, Thiago F. Cordeiro and Tainá S. Araujo for helping with data collection and animal care; Marilia F. Erickson, Larissa M.O. Prado and Vinícius G.S. Santana for contributing with reflectance measurements and visual modeling; and Louise M.O. Prado for figure construction.

